# Machine Learning Method for Optimizing Coding Sequences in Mammalian Cells

**DOI:** 10.64898/2026.01.26.701778

**Authors:** Elias Theodorou, Michael Stadler, Claes Gustafsson, Mark Welch

**Affiliations:** ATUM, 37950 Central Ct, Newark, CA 94560; Department of Genetics, Stanford University, CA 94305

**Author notes:** Communicating authors (Claes Gustafsson, Mark Welch).

## Abstract

Mammalian cell lines are the preferred hosts for producing commercially relevant therapeutic proteins such as antibodies, multispecifics, and cytokine fusion proteins. Even though significant investment is made to optimize upstream and downstream processes, the optimal gene design parameters for heterologous recombinant protein expression remain poorly understood. We describe here a generic approach to gene optimization in which design features are systematically sampled and modulated iteratively using machine learning (ML). Synthetic genes encoding the Dasher fluorescent protein, differing only in synonymous codons, were used to interrogate the gene-sequence preferences of transient antibody-expressing HEK293 cells. Synonymous codon variations influenced expression by more than two orders of magnitude. This variation in protein yield was used to build ML models relating gene design features, which were then employed to design further-improved genes. The ML models were shown to be expression system-specific. Messenger RNA levels and ribosome occupancy were highly correlated with protein levels, suggesting that mRNA lifetime has a causal relationship with coding bias. Our results illustrate a novel, generally applicable method to improve gene expression via synonymous re-coding for any protein target or host cell.

## INTRODUCTION

Recombinant protein expression is fundamental to all biotechnology. Achieving sufficiently high protein yield is critical for reducing the cost of manufacturing biopharmaceuticals and for advancing promising commercial candidates from discovery to development. Even at the discovery stage, biochemical and structural studies are frequently hampered or rendered impossible by insufficient yields from research-scale recombinant protein expression systems. Protein expression yield is considered a function of protein sequence, structure, stability, activity, and interactions with the host. There is also ample evidence that the open reading frame (ORF) coding sequence can greatly influence expression, even though the multitude of underlying sequence design factors remains poorly understood (Gustafsson *et al*., 2004). Importantly, in many biotechnological applications, there can be considerable value in even modest yield improvements. However, currently identified design parameters and associated models are insufficient to predict expression. We sought here a means to reliably maximize gene performance across any host system, independent of a mechanistic model of heterologous recombinant protein expression.

The use of ML methodology combined with classic Design of Experiment (DoE) is well established across engineering disciplines. DoE is common among specific biotechnology applications such as fermentation and process optimization. In a typical study, a range of parameters are systematically varied across a test sample set, and the results are used to build a statistical model that identifies and relates critical features to system performance. For example, one may sample process factors such as temperature, dissolved oxygen, and pH as features and model their relationships with protein expression yield during fermentation. Robust and accurate models help reveal system limitations, enabling engineers to improve quality, throughput, and efficiency while mitigating risk and reducing costs.

In addition to the rich understanding of the environmental factors that affect expression, investments in optimizing the genetic components of expression systems are also likely to yield great success. DoE/ML methodology has, *e.g*., been used to engineer the protein sequence of commercially relevant enzymes to improve enzymatic fitness (Liao *et al*., 2007; Govindarajan *et al*., 2015) and to engineer metabolic pathways for the manufacturing of natural products (Young *et al*., 2018). Here, we explore whether mammalian gene design parameters can be systematically sampled and whether engineering optimization methods can be applied at the gene level, as they have been used in fermentation and protein engineering studies. Current advances in synthetic biology enable the rapid generation of any gene-coding sequence, which can be paired with modular expression systems, empirically validated, and built upon machine learning algorithms to make the Lab in the AI Loop.

Many design features within a gene’s open reading frame have been shown to affect its expression, and these are usually interdependent. The relative frequencies of different codons used to represent each amino acid, codon-pair context, the propensity of the N-terminal end of the mRNA to fold into initiation-competitive structure, the fraction of the mRNA composed of G and C bases, runs of homopolymers, and many other features, have all been proposed as significant determinants of a gene’s expression (Welch *et al*., 2009a, Gustafsson *et al*., 2012). Interdependencies, such as intrinsic correlation between total GC content and secondary mRNA structures, further complicate attempts to provide mechanistic explanations for observed expression levels.

In the past, researchers have typically either compared naturally occurring sequences, made large semi-random libraries, or made a limited hypothesis-driven test gene set. Consequently, evidence supporting any hypothesis has been sparse and/or biased. Instead, we approach the problem from an agnostic perspective, using systematically varied data points and ML-driven inductive reasoning (Gustafsson *et al*., 2016).

With current gene synthesis technologies, we can control any defined sequence variable, enabling systematic and unbiased interrogation of gene sequence preferences. Over the past few years, researchers have begun to exploit this capability to gain a deeper understanding of how gene sequence influences productivity and regulation. Previous work on the gene sequence preferences of *E. coli* showed that gene ranking for heterologous overexpression was poorly explained by host codon bias even in the absence of strong predicted mRNA secondary structure in the translation initiation region (TIR) (Welch *et al*., 2009b). This contradicts the previous dogma that codon frequency, reflected mainly in cognate tRNA levels, should correlate with translation elongation step rates, and that those step rates in turn should govern protein expression yield. As more sophisticated studies have been conducted using current bioengineering tools, we are getting a much more complex and fascinating picture of the many ways in which sequence and host influence expression (Boël *et al*., 2016; Frumkin *et al*., 2018; Rodriguez *et al*., 2024).

Prior work interrogating gene preferences has almost exclusively focused on bacterial hosts, but there is significant commercial value in understanding the preferences of industrially relevant mammalian production hosts. In this work, we demonstrate a method to interrogate the gene-sequence preferences of mammalian HEK293 cells for transient expression. We combine information-rich gene sequence design and systematic testing with machine learning to develop improved gene design algorithms. The strategy does not require prior knowledge or assumptions about genetic preferences or mechanisms. It provides a custom-designed algorithm for the expression system from which the empirical data are derived. This ML-driven approach can be applied to any host, expression condition, or target protein species.

## RESULTS

### Broad Sampling of Coding Space Using a DoE-Designed Gene Variant Test Set

For our initial interrogation of synonymous coding preferences, we designed 48 genes encoding Dasher GFP (Duran *et al*., 2013) to broadly sample codon usage across all amino acids coded by two or more codons (Fig 1). All-against-all sequence pairwise similarity across the set was kept minimal (∼72%) using a combination of Monte Carlo methods and Genetic Algorithms to maximize diversity and minimize co-variation between codon usage parameters and local sequence features. Cryptic splice sites (Kapustin *et al*., 2011) and other previously characterized motifs that have been suggested to be predictably deleterious were avoided in all designs. We also avoided predicted sequences encoding secondary RNA structure for the segment from the Kozak sequence through the first 50 bases of the ORF (ΔG <-10kcal/mol). Frequencies for each of the 59 sense codons (not counting single codons encoding amino acids methionine, tryptophan, or the three stop codons) varied by at least 2-fold, and covariation between usage frequencies for non-synonymous codons was minimized. Our goal was to maximize the information content of the set by minimizing covariation among potentially influential features. In this way, we aimed to avoid confounding effects arising from a hypothesis-constrained design.

**Figure 1.**
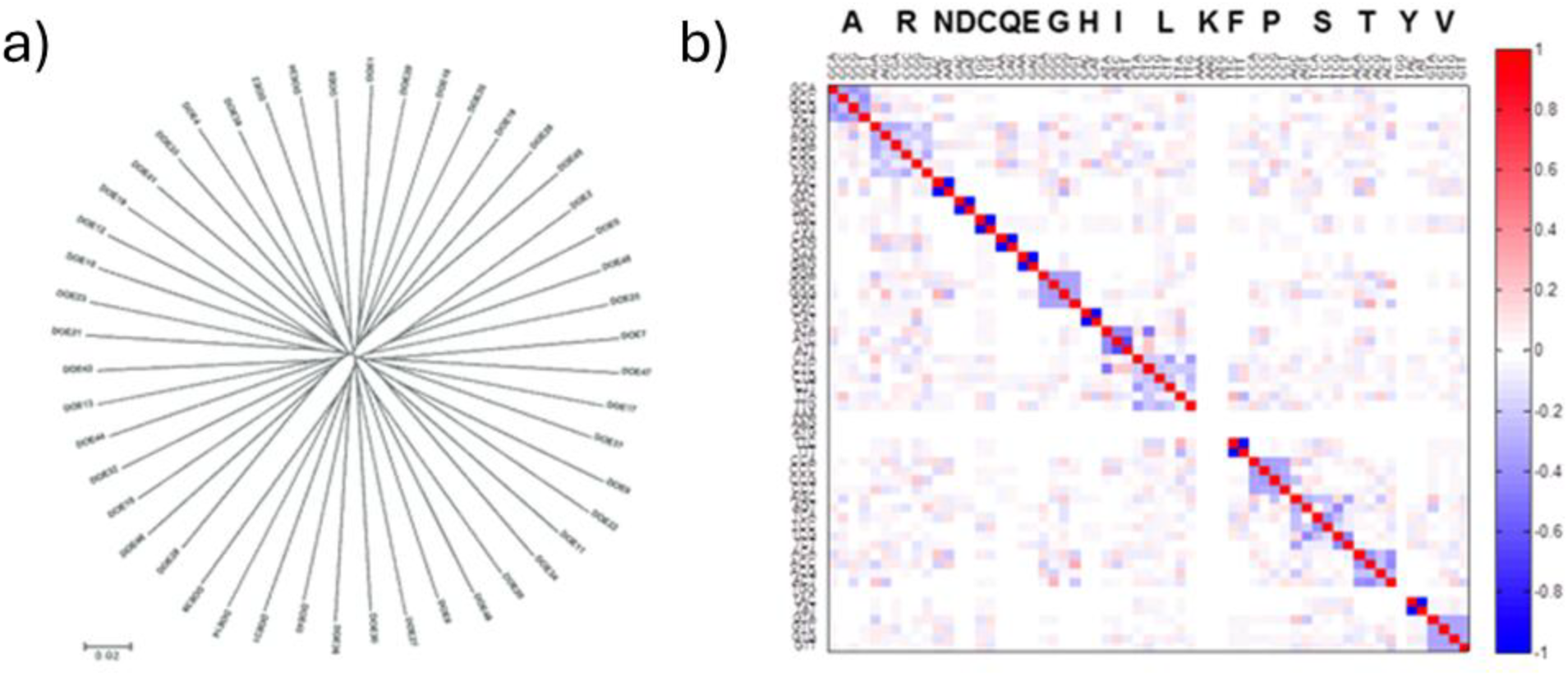
The 48 Dasher genes were designed using Design of Experiments principles to ensure an even, information-rich, and systematic exploration of all coding features without altering the amino acid sequence of the Dasher protein. 1a) An alignment of the designed Dasher genes was plotted as a homology-based neighbor-joining tree generated in MEGA7 (Kumar *et al* 2016). The MEGA7 tree display indicates that each gene is equidistant (∼72% identity) from all other genes in the set as defined by global homology. 1b) Codon usage correlation for the synthetic Dasher sequences. Colors indicate the degree to which each pair of codons is correlated in global usage frequency, either positively (red) or negatively (blue), among genes of the data set. Correlation plots were generated using the PLS Toolbox (Eigenvector, Inc.) in MATLAB. The correlation plot illustrates that the global correlation for all-against-all codons is minimal.

### Impact of Synonymous Changes on Expression Level

Each of the 48 designed test genes was cloned downstream of a CMV promoter in vector pJ609 (catalog vector available from ATUM). The vector is a typical transient vector with a pUC origin of replication and ampicillin resistance for *E. coli* amplification, SV40 origin of replication and puromycin resistance for mammalian amplification. Each vector construct was used to transfect HEK293 cells using standard procedures. Dasher expression was measured by FACS three days after transfection. Expression level, measured as mean fluorescence per cell, ranged over two orders of magnitude across the test set. Previous studies have confirmed that Dasher/GFP fluorescence in *E. coli*, as measured by fluorimeter plate readers, is directly correlated with protein expression yield as quantified by band intensity on polyacrylamide gel electrophoresis (Kudla *et al*., *2009*; *Welch et al*., 2009b). We show that expression variance does not significantly correlate with any of the following: Relative Synonymous Codon Usage (RSCU; Ikemura 1981), Codon Adaptation Index (CAI; Sharp *et al*. 1987), predicted mRNA structure strength at the translation initiation region, GC content, or CpG prevalence. Of these commonly used criteria for gene optimization, none showed a statistically significant correlation with expression yield (data not shown). This study indicates that a simple frequency match between the introduced gene and the host’s implied tRNA pool is insufficient to predict—let alone maximize—performance.

### Modeling the Impacts of Coding Features

In a previous 2009 paper, we used Partial Least Squares (PLS) regression to identify and quantify relationships between gene features and expression in *E. coli*, analyzing a broad range of sequence features to construct a model that can be used to drive synthetic gene design (Welch *et al*., 2009b). For the study presented here, we instead applied a suite of AI/ML algorithms to capture the correlation between sequence design features and recombinant protein expression in transient HEK293 mammalian cells. We obtained models with an R^2^(CV) of 6.2 (Fig. 2, first-round blue datapoints). Interestingly, most of the variation in expression yield is predicted by codon usage frequency features. To note, the human codon bias is predicted to be suboptimal for expression in the transient HEK293 system.

**Figure 2.**
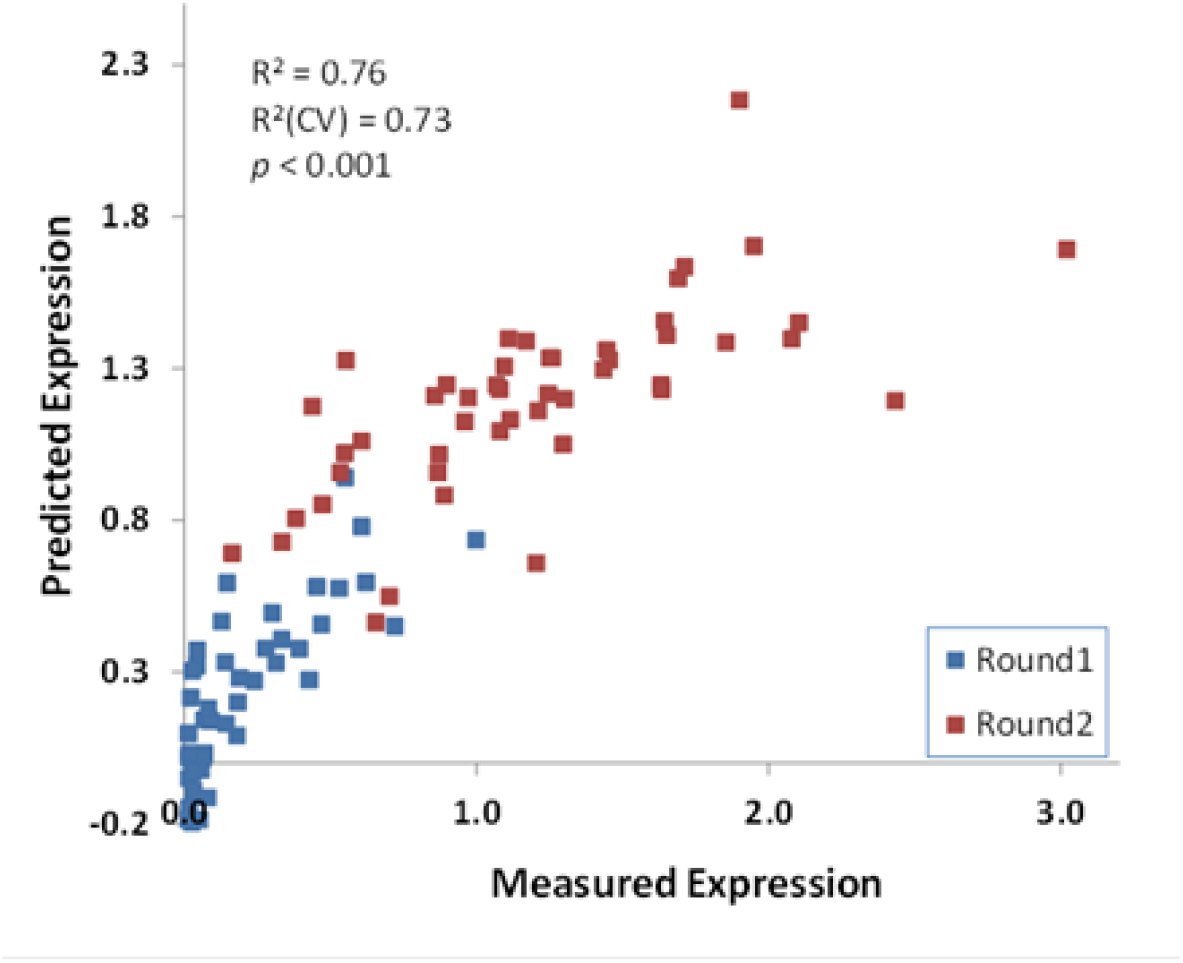
Empirical gene optimization for Dasher expression in transient HEK293. Protein expression of the Dasher enzyme is measured by flow cytometry, with excitation at 505nm and emission at 525nm, and the measured values are normalized to the highest expressing variant on the X-axis. The model fit for the expression level as a function of codon usage frequencies is shown. For each gene variant, the predicted expression level (Y-axis) was plotted against the measured expression level (X-axis). Expression predictions are shown for the agnostic round 1 (blue) and the ML model-improved round 2 (red). R^2^ (CV) indicates the correlation coefficient for the fit of the full model in cross-validation bootstrap, where 20% of the data are randomly left out of the regression.

**Figure 3.**
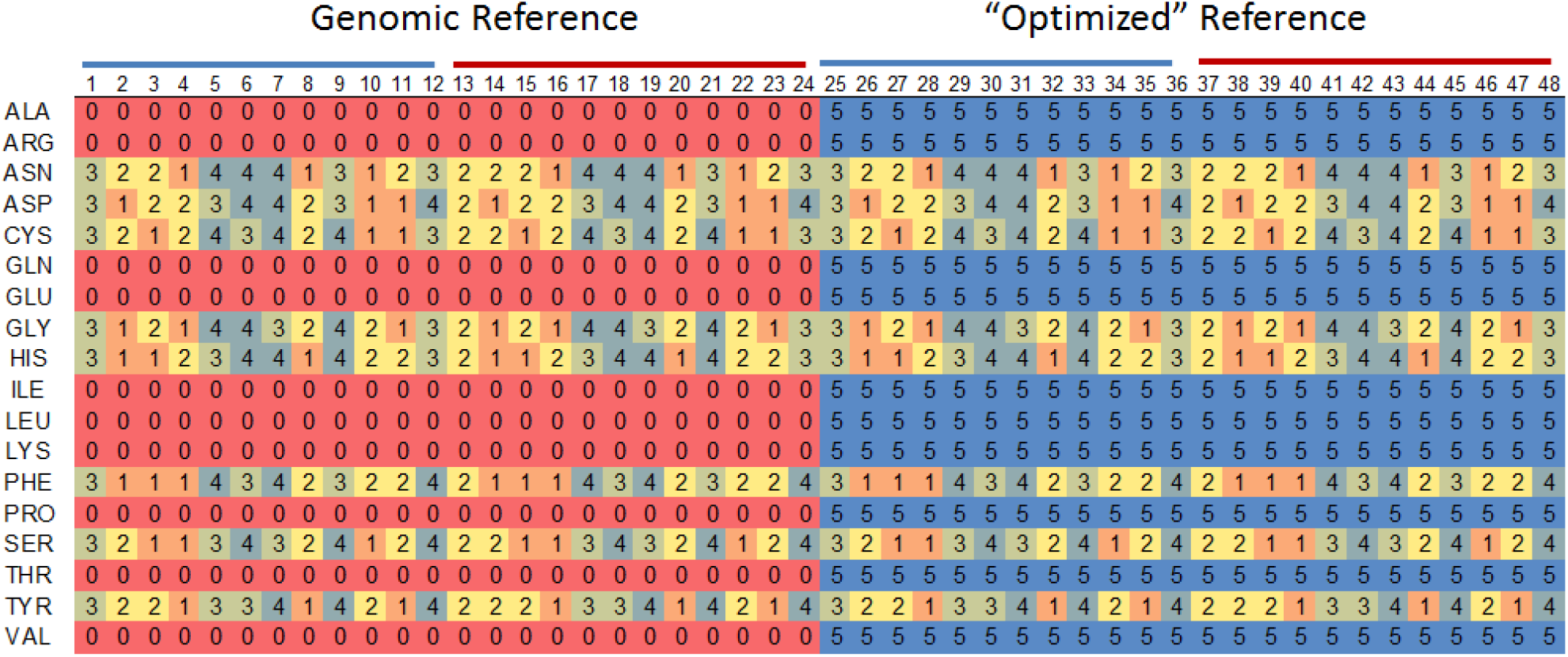
Design of the matrix dataset. Half the dataset (24 Dasher genes, columns 1-24) was designed to capture the contribution of G:U wobble codons in the genomic coding background. The other half (columns 25-48) captures the contribution of G:U wobble codons in the optimized round 2 background. Each color and number in the matrix is a non-descript label corresponding to a predetermined codon bias for the amino acid in the first column. Thus, 0 on a red background (e.g., Gln) defines a specific ratio between the codons CAA and CAG in the genomic reference sequence. Number 5 on blue similarly illustrates the corresponding ratio of Gln codons CAA and CAG in the optimized reference sequence. The wobble incorporation is identical across, e.g., gene variants 1, 25, and so on.

**Figure 4.**
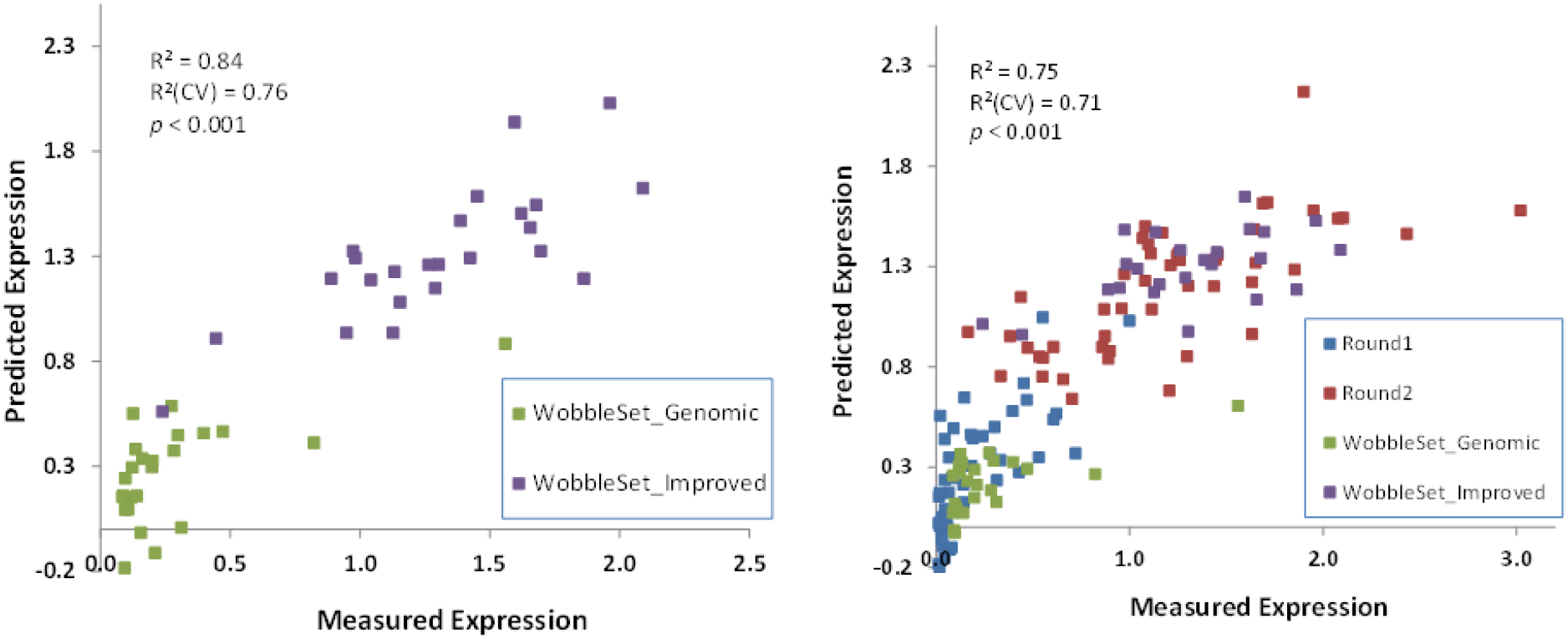
Gene optimization exploring the contribution of G:U wobble in the context of gene variants designed using genomic (green) or optimized (purple) non-wobble codons. The model fit of expression level as a function of codon usage frequencies is shown. For each gene variant, the measured expression level was plotted against the predicted expression. The R^2^ (CV) is the cross-validated correlation coefficient for the model, with 20% of the data held out. The a) panel presents results showing how contributions of wobble codons are assessed in the context of genomic codon bias (columns 1-24 in Figure 3) and in the context of optimized codon bias (columns 25-48 in Figure 3). The b) panel combines data from Fig. 4a and Fig. 2 to form an integrated model.

### Model-Driven Design for Improved Expression of Genes

Sampling all possible ORF coding sequence variation is impractical due to the near-infinite number of possible sequences. In our approach, we use diversification via systematic variation and ML to point toward the variable space where better performance is observed. We intentionally do not invoke any mechanisms in design or modeling to avoid biasing the interpretation of the results. Our goal was to develop a method that could be applied, independent of mechanistic knowledge, to any protein target or expression system. With iterations of diversification and modeling, we should be able to walk in sequence-expression-space toward more optimal coding, even if the new predicted sequences are outside the training set. Accordingly, we modeled gene sequences from the original training set towards a second round of improvement.

We first used the model to define a set of sequence features reliably predicted to be superior to the first-round variants. This new reference was then used as a starting point for variation, using the same methodology as in the first round, except that variation was emphasized for codons identified as most predictive in the first round. Forty-eight new Dasher genes were designed, cloned, and again tested for transient expression in HEK293 cells, as in the first-round set. As in the first round, a wide range of expression levels was observed, and both the average and best performance shifted towards higher Dasher expression yields, demonstrating the first-round model’s ability to predict improved gene design beyond the test set. This second round of data was used to augment the first, and a combined model was obtained, which served as a guide for subsequent gene design (Fig. 2).

### Impact of Wobble Codons

G:U wobble codons decoded by tRNAs using non-canonical G:U base-pairing at the third codon position have been suggested to be associated with an increase in ribosomal pausing in higher eukaryotes, implied to result in lower protein expression yield (Stadler *et al* 2011). We accordingly decided to examine the preferences and effects of wobble codons on heterologous protein expression in our Dasher test system. Using a similar strategy as described above, we devised a 48-gene test set that systematically varied wobble codon usage for the eight amino acids (C, D, F, G, H, N, S, and Y) that have G:U wobble codon options. For 24 of the designed genes, the 10 non-wobble amino acids were encoded using human average codon usage frequencies derived from the Kazusa database (Nakamura *et al* 2000). For the second 24 genes, the ten amino acids were encoded using a codon frequency predicted to be improved relative to the initial interrogation described in Fig. 2 (the same as the reference frequencies used for the second round of interrogation). Predicted deleterious motifs were avoided in all designs, and codon assignment by position was randomized to maximize sequence diversity at wobble and non-wobble amino acid positions.

Our results suggest that a majority of the variance is explained by coding for the ten non-wobble amino acids. A significant 4.8-fold shift in average expression level was observed for the subset using “improved” codon usage for the non-wobble amino acids. Wobble codon usage alone did not explain any significant variance in protein expression yield in this dataset.

### mRNA Level is Strongly Correlated to Expression

To gain insight into the mechanisms underlying the observed relationship between protein expression and coding preferences, we analyzed total mRNA and ribosome occupancy for a representative subset of gene variants using sequence-based ribosomal profiling (Ingolia *et al* 2009). The total specific mRNA level, as measured by RNA-seq, correlates well with the observed protein expression level (Fig. 5a).

**Figure 5.**
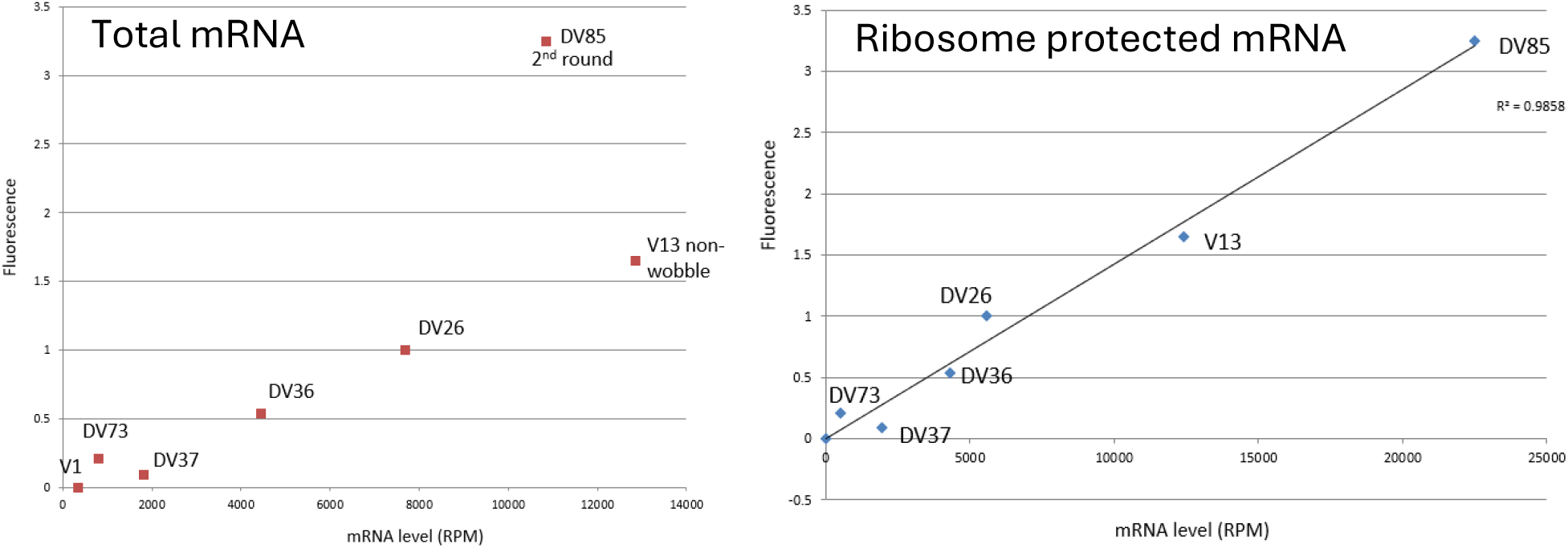
a) Total Dasher mRNA abundance vs. protein activity. Protein activity (Y-axis) is measured as fluorescence, normalized to that of the Dasher variant DV26. Dasher mRNA level (X-axis) is quantified by RNA-seq and reported as reads per million (RPM). b) Ribosome-protected Dasher fragment abundance vs. protein activity. Protein activity (Y-axis) is measured as fluorescence normalized to that of the Dasher variant DV26. Ribosome-protected Dasher mRNA fragments (X-axis) are quantified by RNA-seq and reported as reads per million (RPM).

An even stronger correlation with fluorescence was observed for the number of ribosome-protected fragments (RPFs), illustrated in Fig. 5b. A comparison between mRNA levels and RPF suggests that the most highly expressed gene, DV85, has increased ribosome occupancy per mRNA. This variant shows ∼40% higher protected mRNA fragment levels than V26 (the best first-round variant), yet >3x the expression. It is worth noting that the Dasher gene variant DV85 is the only variant from round two of the optimization assessed for RPF.

The strong correlation observed between protein fluorescence and total mRNA levels suggests that the primary mechanism by which optimal synonymous recoding enhances protein yield is not necessarily faster translation (elongation velocity), but instead increased steady-state mRNA abundance or functional lifetime. However, the single datapoint from the second-round DV85 is an outlier, producing twice as much protein per total mRNA. We interpret this to imply that the increase in protein expression from round one to two is primarily driven by more efficient ribosome loading.

### Synonymous Coding Effects are Dependent on the Expression System

To address differences in gene preferences across hosts, we also tested the same initial 48 variant Dasher ORF sequences for expression in *E. coli* and in baculovirus Sf9. Each ORF sequence was cloned into expression vectors for each host, and the Dasher protein was expressed under standard conditions (Jarvis DL, 2009). As with HEK293, we observed a wide range of protein expression in these hosts; however, gene preferences differed between expression systems. No pair of the three expression systems showed correlated gene rankings (Fig. 6). Interestingly, genes expressing near the high end of the distribution across all expression systems could be identified. As with HEK293 expression, neither natural codon bias in the host nor the predicted RNA lowest-energy structure in the translation initiation regions explained protein expression yield. Host-specific predictive gene design algorithms were built, as described above, for *E. coli* and the baculovirus Sf9 using 48 systematically varied gene variants (data not shown). The *E. coli* algorithm derived from the 48 Dasher dataset presented here bears no similarity to either the commonly used CAI or RSCU gene-sequence optimization algorithms for *E. coli*. However, it is strikingly similar to the *E. coli* algorithm described earlier (Welch *et al*., 2009b).

**Figure 6.**
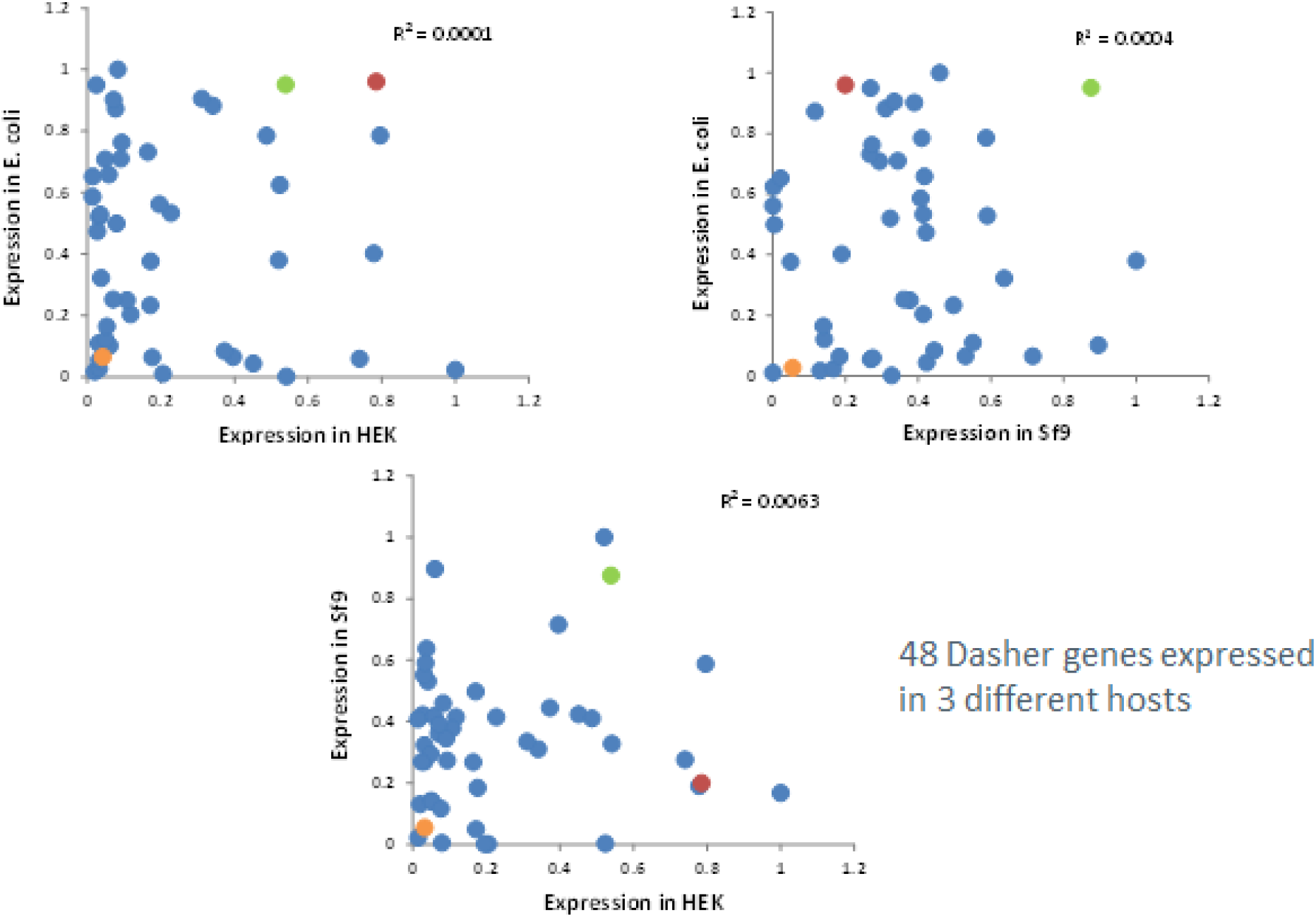
Protein expression of the same 48 Dasher genes in *E. coli*, HEK293, and Sf9. Expression is normalized to the highest-expressing gene variant in each host. Even though there is no apparent correlation in coding preference between the hosts, there are examples of Dasher coding gene variants being well expressed in all dimensions (green), poorly expressed in all dimensions (orange), and well expressed in two hosts and poorly expressed in a third (red).

### Synonymous Coding Effects are Protein Independent

The value of a codon optimization algorithm is directly correlated with its level of generality. If each protein requires a new algorithm built from 48 independent genes and measurements, the algorithm’s value is limited from a bioengineering perspective. To address this concern, we designed ∼50 gene variants for a representative subset of six recombinant proteins, totaling 302 synthetic genes. Each gene sequence was designed and synthesized as outlined above in the Dasher example to minimize any inherent variable correlation. Expression titer in transient HEK293 cells was determined by polyacrylamide gel electrophoresis for each of the gene variants, and the titer was normalized to 1 for the highest expressing variant for each protein. The predicted expression titers for each gene variant were calculated using the algorithm derived from the Dasher dataset (above), resulting in a reasonable (R^2^ CV=0.64) and consistent prediction of relative expression yield (Fig. 7). The Dasher-derived algorithm predicts the relative expression titers well. However, the absolute expression titers between each protein vary by more than an order of magnitude. We infer that this discrepancy is due to inherent differences in protein stability, folding efficiency, and other protein-specific factors.

**Figure 7.**
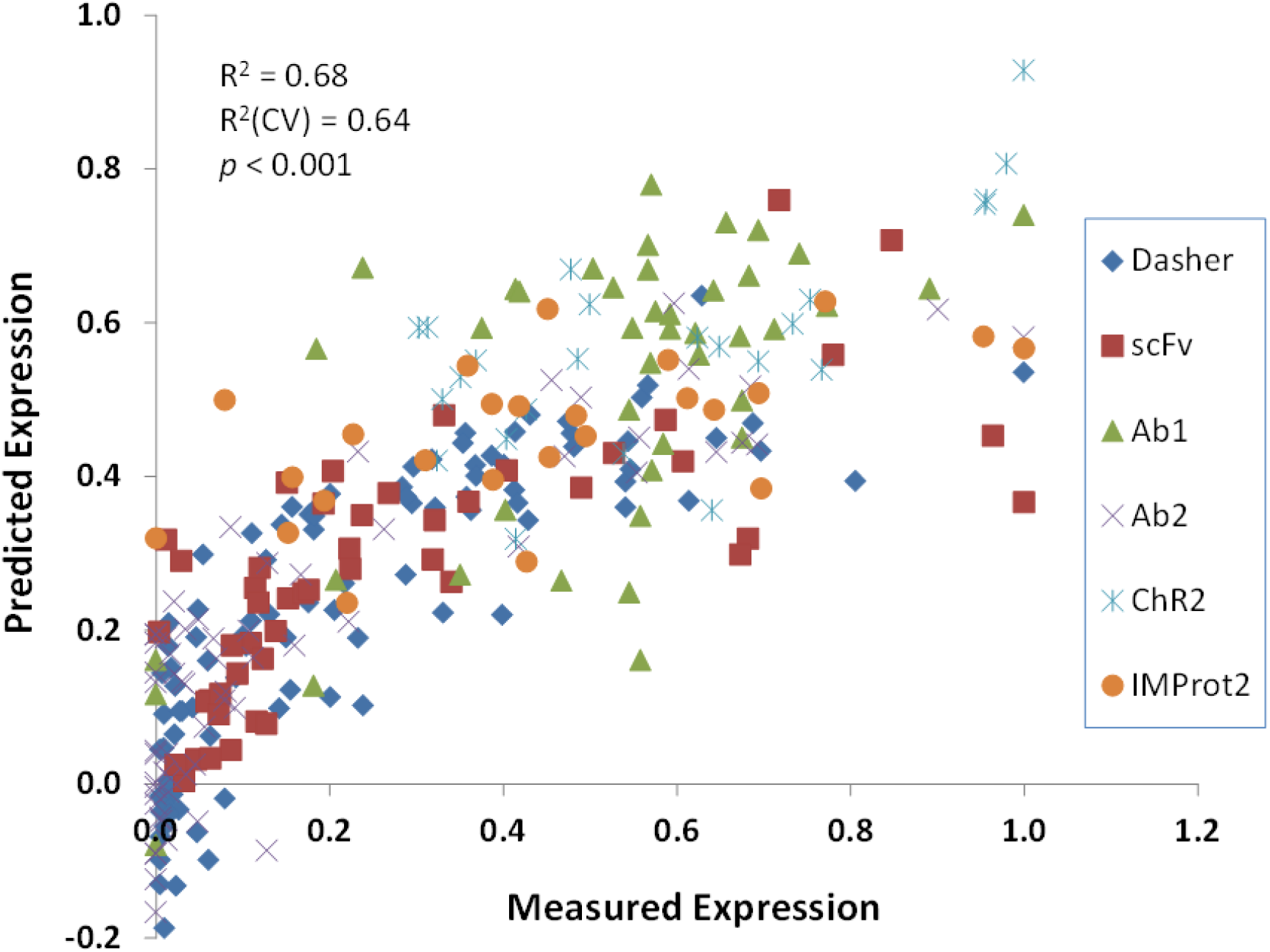
Protein expression of 302 genes encoding six different proteins: Dasher (fluorescent protein), scFv (single chain variable fragment), antibody #1 and #2, and membrane proteins ChR2 and IMProt2. Each corresponding gene set (typically 48 gene variants encoding each protein) is designed using the systematic variance approach described above. Each gene sequence variable is as non-correlated as possible to any other variable in the same set (see Fig. 1a and 1b for illustration of the Dasher set) while not altering the protein sequence. Genes are inserted into transient expression vectors and transfected into HEK293 cells using standard procedures. Recombinant protein expression after 3-5 days was quantified by BLI (antibody #1 and #2), PAGE, or FACS (Dasher), and the highest expressing protein from each set was normalized to 1 on the X (Measured Expression) axis. The predicted value on the Y axis was quantified from the algorithm that was built solely on the Dasher data (Fig. 4b). Despite the heterogeneity of the protein structures and compositions, the sequence origins and expression yield, we see a significant consistency in the expression data (p< 0.001, R^2^(CV) = 0.64) suggesting that coding effects are essentially protein independent within the same host.

## DISCUSSION

Empirical optimization of gene design using ML represents a paradigm shift in heterologous protein expression. By systematically exploring the vast search space of synonymous codon variants and correlating sequence features directly with protein yield in a host-specific manner, our findings dismantle long-held assumptions and establish a novel, universally applicable strategy for the efficient expression of recombinant proteins. Not only does the methodology described here provide a foundation for engineering high protein expression yields, but it also offers a convenient format for mechanistic studies, e.g., whether wobble codons affect protein expression (they do not) or whether coding affects mRNA stability (it does).

Most existing codon-optimization models are based on the non-uniform codon distribution of naturally occurring sequences. In natural evolution, overproducing any protein is a waste of energy and results in rapid extinction from the gene pool. We further see no significant correlation between natural codon bias and recombinant protein expression yield. Other codon optimization algorithms are based on a small number of extreme structure predictions vs a non-structured parental sequence (Zhang *et al*., 2023). An alternative approach is to use modified nucleotides in the mRNA to alter its structure (Mauger *et al*., 2019), a method not directly applicable to recombinant protein expression.

The functional outcome of data modeling is a direct reflection of the information geometry of its training data. To achieve a model with a predictable outcome, the data distribution must not only be large in quantity but also representative and evenly distributed across space. In the context of modeling sequence-function data, the uniform distribution of data points within a multi-dimensional feature space is the primary determinant of the model’s accuracy. If an AI algorithm is trained on a cluster of data points that occupy only a small corner of the feature space, such as codon bias of natural sequences or highly structured mRNA, it will become highly proficient at making inferences within that cluster. Still, it will likely fail when asked about untested regions of space.

Building a test set of 48 equidistant and systematically varied Dasher genes allows us to model all features, even if we do not know their exact identities. Not only can we generate gene design models that interpolate reasonably well, but the underlying algorithm also extrapolates recombinant protein expression with reasonable accuracy (Fig. 2). Extrapolation is a notoriously hard problem for AI. Utilizing a broad, systematically varied dataset and agnostic feature extraction allows us to overcome this concern.

The methods described here, using unbiased design, testing, and machine learning to improve gene performance, can be applied to genes for any protein expressed in any system, without prior knowledge of the host’s expression biochemistry or regulatory genetics. Furthermore, while the example presented here focuses on codon usage frequency features, the search space also include any variable of interest to improve gene performance further, such as other translation features (mRNA secondary structures, runs of homopolymers, ribosome binding sites etc.), transcription features (promoters, upstream activating sequences, terminators, etc.) and genome features (location, copy number, orientation etc.). Unbiased modeling of the features and their effects on the output not only identifies and quantifies linear additivity in the prediction models but also captures relative epistatic relationships among the identified features.

Several studies in bacterial, yeast, and mammalian cells have shown a significant correlation between protein and mRNA levels for synonymous coding variants (Boël *et al*., 2016, Newman *et al*., 2016, Presnyak *et al*., 2015, Wu *et al*., 2019, Radhakrishnan *et al*., 2016). The correlation between steady-state mRNA levels and protein expression that we observe in this study is consistent with these earlier findings; however, it is not clear by what mechanism the mRNA stability is influenced, and it may be that both mRNA structure and codon usage contribute (Mauger *et al*., 2019).

The ribosome protection experiment suggests that optimal coding sequences, such as those encoded in variant DV85 (the highest-expressing variant), may facilitate more efficient ribosome loading or sustained high ribosome occupancy, thereby protecting the mRNA from ribonuclease degradation. The observation that the best-performing variant showed disproportionately higher RPF relative to its mRNA level suggests that the ML model successfully captured translation efficiency (ribosome occupancy per mRNA molecule) as a critical feature. Our results are consistent with the model of translation-coupled mRNA decay (Doma *et al*., 2006; Shoemaker *et al*., 2012), in which non-optimal coding sequences or ribosome pausing events can trigger mRNA degradation pathways, thereby limiting the time available for translation. The mammalian homologs of Hel2, and through a separate pathway, Syk1 and Smy2 coordinate the regulated decay of translating mRNA as a function of ribosomal states (Veltri *et al*., 2022).

This iterative, ML-driven process outlined here offers a robust engineering solution to what was previously a trial-and-error process. It can be applied to any recombinant protein expression system to build a custom optimization algorithm. The methodology provides a path to reliably achieving maximum expression for commercially relevant therapeutic proteins, accelerating candidates from discovery research to large-scale development. Our approach moves expression engineering from a hypothesis-driven science to a predictive, data-driven engineering discipline, fundamentally changing the timeline and success rate of synthetic biology design.

## Notes

The authors declare the following competing interests: Claes Gustafsson and Mark Welch are co-inventors on granted US Patents that cover the specific method disclosed in this study.

## Acknowledgement

The authors thank Andrew Fire at Stanford University for guidance and funding for the ribosome protection assay disclosed in Fig. 5, Nathan Spangler and Katie Rogers at Aldevron for performing the baculovirus protein expression disclosed in Fig 6, Sridhar Govindarajan, Jeremy Minshull, Tuong Nguyen, and the ATUM team for technical support and scientific discussions. This work was supported by ATUM.

